# Increased attentive use leads to more idiosyncratic functional connections

**DOI:** 10.1101/2025.10.15.682657

**Authors:** Pinar Demirayak, Leland L Fleming, Paul Stewart, Rachel Chua, Kristina M Visscher

## Abstract

Experience is thought to modify neural connections to adapt the network to be more optimal for the environment. Given the brain’s complexity, multiple network changes could each move the system toward optimality. Standard methods ignore this multiplicity and examine each connection independently; these studies have often shown considerable inter-individual variability and modest effects (1). Here, we take a different strategy, determining how a whole-brain connection pattern differs from the typical pattern, that is, how ‘idiosyncratic’ the pattern is. We examined how the idiosyncrasy of whole brain connection patterns varies with frequency of the use of that part of cortex for attention-demanding tasks, focusing on central versus peripheral vision in healthy individuals (where individuals use central vision more frequently for attention-demanding tasks). We found that the whole-brain pattern of functional connections to the cortical representations of central vision is idiosyncratic, whereas patterns of connections to representations of peripheral vision were very similar person-to-person. In a second set of analyses, we examined the brains of people with central vision loss who use a portion of peripheral vision (called the preferred retinal locus) more frequently for attention-demanding tasks in their daily lives. The cortical representation of the preferred retinal locus exhibits more idiosyncratic connections, compared to a control brain region, or compared to the same brain region in matched control participants with healthy vision. These results are consistent with the hypothesis that increased attentive use of a brain area results in idiosyncratic patterns of whole brain connections.

**Significance Statement:** We found that increased attentive use of a brain region results in more idiosyncratic patterns of connections of that region to the rest of the brain. Our findings support the view that V1 retains the capacity for plasticity well beyond the critical period and that these adaptations are idiosyncratic to the individual’s experiences. This approach suggests that tailoring personalized rehabilitation plans for individuals with retinal diseases may be more effective than a ‘one size fits all’ approach. More generally, it offers a promising framework for investigating brain plasticity in both typical and clinical populations, especially in the context of sensory loss and compensatory adaptation.

## Introduction

Comedian Jamie MacDonald, who is blind, said “Some people, they see disability as *one* thing. Blindness has infinite combinations of psychological and physical impacts on people… Blind people, we’re all different. We’re like snowflakes. There are not two of us the same. And if a lot of us fall, people panic” (2). MacDonald’s humorous quote brings home a point that is often ignored when studying brain plasticity: the brain’s response to changes in experience is idiosyncratic. As researchers, we often ignore this point, typically averaging responses among individuals, forgetting that each participant’s experience is like a snowflake.

Inter-individual variability in brain connectivity reflects the interplay between common neural architectures and the unique imprint of individual factors. While the large-scale organization of brain networks is broadly shared across people (3, 4), growing evidence suggests that functional connections also carry idiosyncratic signatures (5, 6). These idiosyncratic features may arise from personal experience or from other factors. Because experience is inherently idiosyncratic, the connectivity changes that accompany it are unlikely to be uniform across individuals. This idiosyncrasy is not evenly distributed across the brain: regions involved in sensory processing, such as the early visual cortex, generally exhibit lower inter-individual variability than higher-order association areas (5, 7). However, the extent to which variability of connection patterns emerges following experience remains poorly understood. To address this, we quantified whether brain regions representing frequently attended portions of the visual field -such as central vision in individuals with healthy sight-show more idiosyncratic versus more stereotyped patterns of whole-brain functional connectivity.

We examined both individuals with healthy vision and those who have lost central vision due to macular degeneration. People with healthy vision preferentially attend to the fovea due to its high spatial resolution (8, 9). Vision subserved by the fovea and macula of the retina is critical for reading, recognizing faces, and scrutinizing whatever objects or scenes a person attends. Occipital and frontoparietal attention networks are more active when stimuli are presented foveally (as opposed to peripherally), reflecting a natural bias of attentional systems toward central vision (10). Thus, the brain regions associated with central vision habitually undergo more attentive use than those associated with peripheral vision.

A severe scotoma in central vision produces a profound and enduring disruption of visual input to the fovea, with severe consequences for everyday life (11). Central vision loss due to macular degeneration (MD) damages the macula while sparing much of the peripheral retina (12). Individuals with MD often adopt a preferred retinal locus (PRL) in the intact peripheral retina as a new fixation point for their daily attended activities. The PRL can support tasks that are normally mediated by the fovea, such as scrutinizing objects and word recognition, but at reduced spatial resolution and with altered oculomotor dynamics (13, 14). Visual processing of information in the PRL thus changes dramatically, and habitual attention shifts from the fovea to the PRL. Due to the retinotopic organization of the early visual areas in humans, central vision loss is an ideal model to study experience-dependent changes in visual processing.

Resting-state functional connectivity reveals both shared and individual-specific features of brain organization, reflecting the balance between canonical large-scale networks and idiosyncratic connection patterns. Connections within networks, such as the default mode network, the dorsal attention network, and sensorimotor networks, show strikingly consistent strong connections in almost all individuals (6). Yet alongside this stereotypy, there are unique patterns of fine-grained functional connections (6, 15). Fine-grained patterns of connectivity vary substantially between people, capturing the influence of individual anatomy, development, and experience (5, 16), resulting in patterns of connections that are unique to an individual and are sometimes called “connectome fingerprints.” Building on this framework, we asked whether the functional connections of visual field representations-particularly those habitually engaged for attention-also carry such individualized signatures, and whether these signatures change in the context of central vision loss.

We examined how different features of sensory experience (i.e. sensory deprivation and increased usage) affect functional connectivity of the cortical regions corresponding to retinal lesions and areas of increased usage on the cortex. Our data are consistent with the hypothesis that brain regions that experience ‘more attentive use’ show more idiosyncratic patterns of whole-brain connections.

## Results

### Central vision functional connectivity (FC) maps are more idiosyncratic than peripheral vision FC maps in healthy vision participants

To examine the individual variability of functional connectivity patterns associated with different parts of the visual field, we compared the spatial similarity of FC maps derived from central and peripheral visual representations for participants with healthy vision. We calculated the idiosyncrasy based on similarity of an individual’s functional connectivity map to a group of healthy control participants, as described in Figure 1.

**Figure 1.**
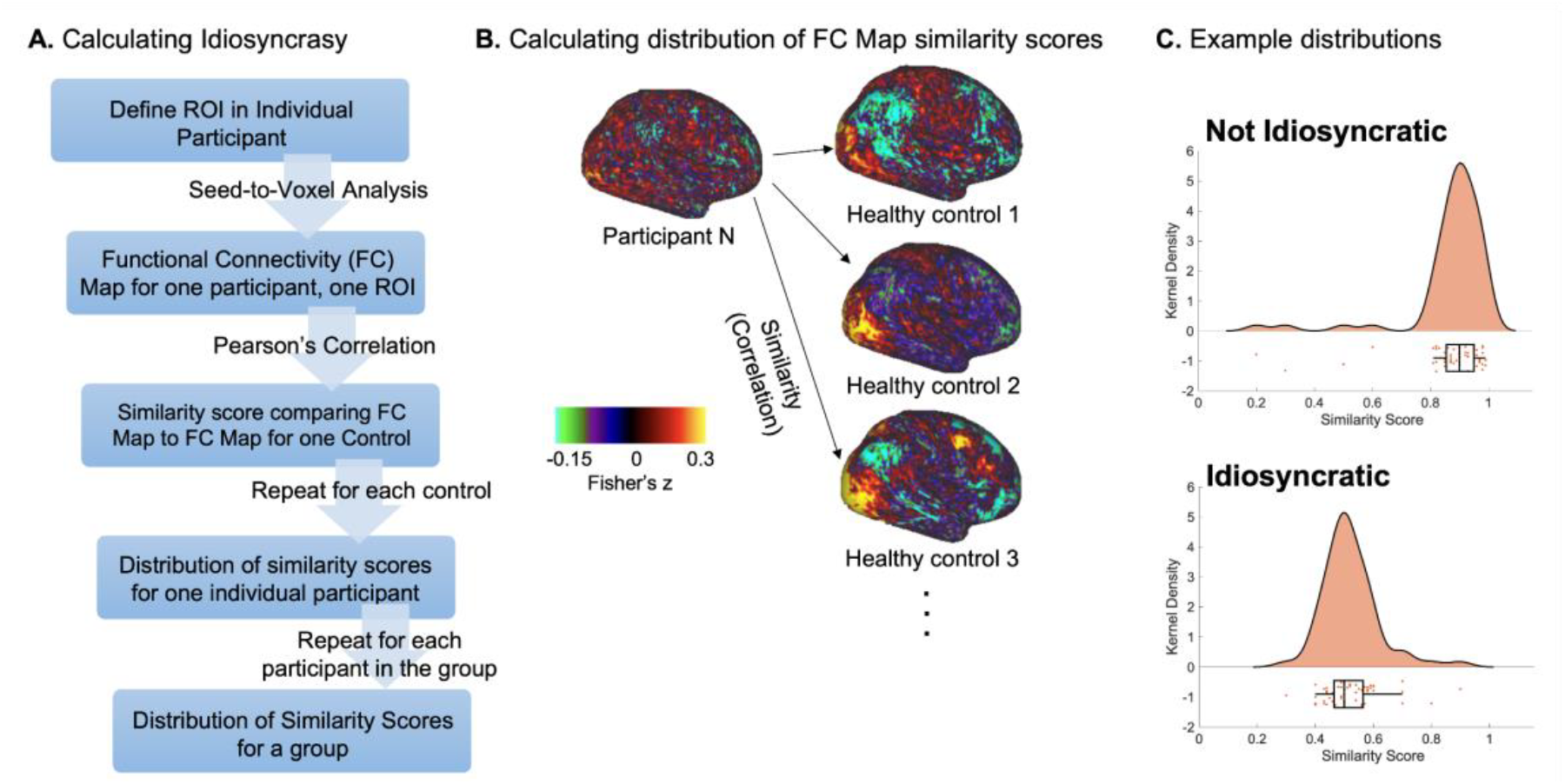
**(A) Schematic representation of how a distribution of similarity scores is calculated. (B) Graphical description of the ‘Distribution of similarity scores for one Individual participant’ step in A.** For each region of interest, for each participant, we calculate the similarity between that participant’s FC map and the corresponding FC map for every healthy control. **(C) Example similarity histograms** for a group with more stereotypical, non-idiosyncratic patterns of connections (top), and more idiosyncratic patterns of connections (bottom).

Our results revealed that FC maps associated with central vision exhibited significantly lower inter-subject similarity compared to those derived from peripheral vision, indicating greater idiosyncrasy in the central visual field’s connectivity profile (Figure 2). ANOVA showed that the main effect of Region of Interest (ROI, central vs. peripheral) was significant for the connectivity FC maps of central and peripheral vision processing regions within healthy vision controls (F(1,10877)=8.5697 p=0.0034). Our results suggest that more attentive usage of central vision in individuals with healthy vision is associated with more idiosyncratic FC maps of central vision representations than peripheral vision representations.

**Figure 2.**
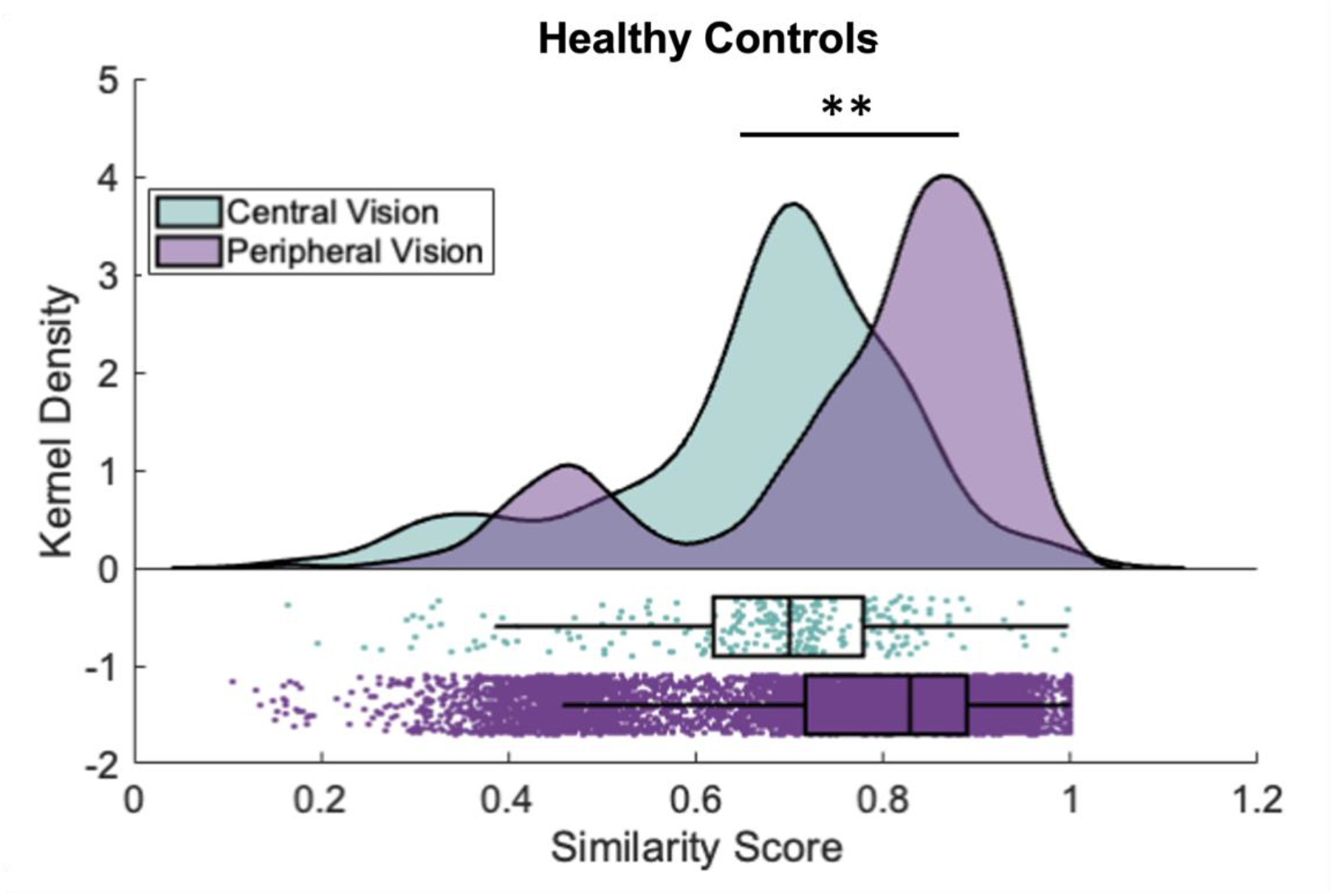
Distribution of similarity scores for FC maps seeded with cortical representations of central and peripheral vision in healthy vision individuals. Note that the maps for the central region of interest (green) are significantly more idiosyncratic than the peripheral region of interest (purple) for healthy vision controls. Colored data points indicate individual measurements. Histograms show the frequency distribution.

### Experience with loss of attentive use is associated with no change in overall idiosyncrasy

We also examined idiosyncrasy of functional connection patterns comparing participants with central vision loss due to macular degeneration (MD) to healthy controls (HC). Participants with central vision loss had lost retinal input to the center of vision; the Lesion Projection Zone (LPZ) is the associated portion of cortex. A control peripheral vision processing region was defined in all participants, where retinal input was similar to the PRL, but the area was not used habitually for attentive tasks; this region is called the Unpreferred Retinal Locus (URL) and the associated portion of primary visual cortex is called the cURL. We compared FC maps’ similarity between LPZ and cURL in MD and HC participants by implementing linear mixed effect model ANOVA (Figure 3). We found a statistically significant main effect of ROI (F(1,6528)=10.017, p=0.0016) and interaction between ROI (LPZ, cURL) and diagnosis of the participants (MD vs HC) (F(1,6528)=16.849, p<0.001). When we directly compare similarity scores from central vision processing region (LPZ) and peripheral vision processing region (cURL) in V1 between the individuals with central vision loss and individuals with healthy vision by using an independent sample t-test, (Figure 3C) similarity scores based on both LPZ (t(734)=0.9899, p=0.32) and cURL (t(5794)=-1.757, p=0.079) are not different between groups (Figure 3C). Mean (μ) and standard deviations (σ) of similarity scores of MD patients vs healthy vision controls based on ROIs were: MDμ_LPZ_:0.6592, MDσ_LPZ_:0.1887, HCμ_LPZ_:0.6728, HCσ_LPZ_:0.1534, MDμ_cURL_:0.7908, MDσ_cURL_:0.1839 and HCμ_cURL_:0.7768, HCσ_cURL_:0.1662.

**Figure 3.**
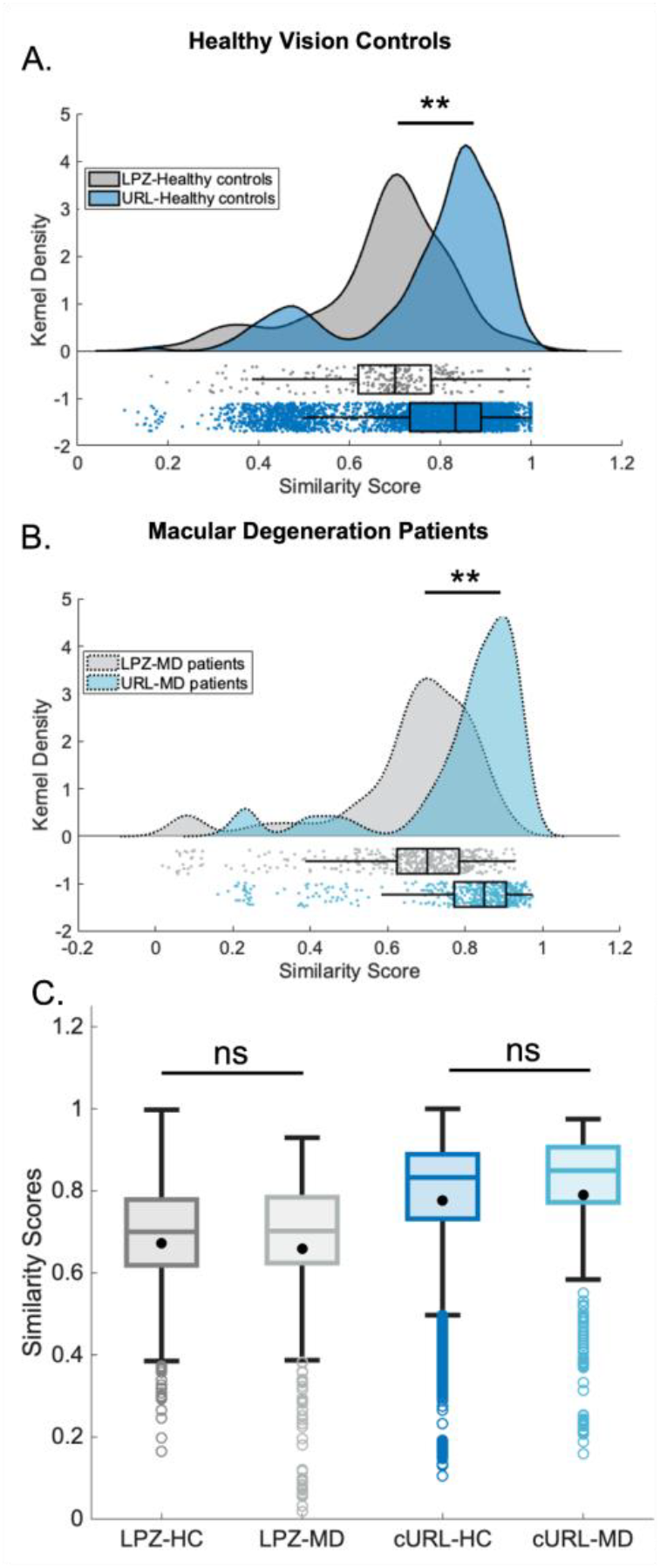
Central vision processing cortical region (LPZ)-seeded functional connections are more idiosyncratic than cURL-seeded functional connections in both macular degeneration patients and healthy vision controls. Distribution of similarity scores of LPZ- and cURL-seeded FC maps in healthy vision controls (A) and macular degeneration patients (B). (C) Group-level independent sample t-test comparisons of similarity scores did not find a difference between MD and healthy control groups, either for the LPZ- or the cURL-seeded FC maps. The black data point represents the group mean and the colored horizontal line represents the group median. Error bars show ±1 standard error. The top and bottom bounds of each box represent the 75th and 25th percentiles, respectively. The whiskers extend to the minima and maxima data points, not considered outliers. **p<0.01.

### Experience with more attentive use is associated with more idiosyncratic connection patterns in MD patients

We examined idiosyncrasy of functional connection patterns of the cortical region associated with the portion of vision that the MD participants used more, the cPRL, comparing MD to HC. As above, results from this region were compared to the control location, the cURL, where retinal input was similar to the PRL but not used habitually for attentive tasks. We analyzed individual variability in cPRL and cURL-seeded FC patterns to evaluate sensory experience-dependent changes following more attentive usage of cPRL. Distributions of the similarity scores of MD patients and healthy vision controls are shown in Figure 4. We found that the main effect of ROI (F(1,11588)=62.757 p<0.001) and the interactions between ROI and diagnosis of the participants between cPRL and cURL (F(1,11588)=852.56 p<0.001) seeded FC maps are statistically significant. Mean (μ) and standard deviations (σ) of similarity scores of MD patients vs healthy vision controls based on ROIs were: MDμ_cPRL_:0.6456, MDσ_cPRL_:0.1882, HCμ_cPRL_:0.7663, HCσ_cPRL_:0.1695, MDμ_cURL_:0.7908, MDσ_cURL_:0.1839 and HCμ_cURL_:0.7768, HCσ_cURL_:0.1662.

**Figure 4.**
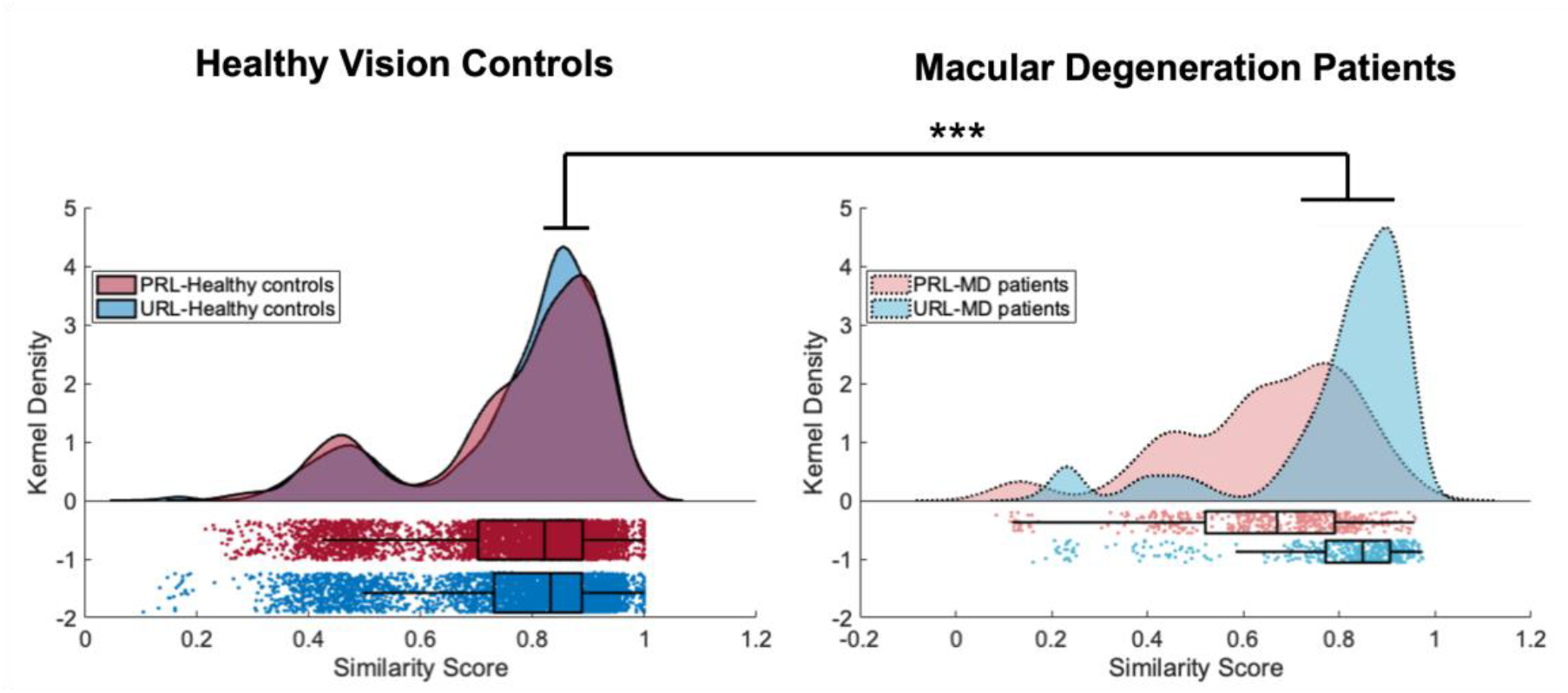
The cortical projection of the preferred retinal locus has more idiosyncratic patterns of FC in MD participants. cPRL (shown in red) and cURL (blue) seeded FC maps’ distribution of similarity scores are shown for macular degeneration patients (dashed line) and healthy vision controls (solid line). Note that more attentive use of a specific region in peripheral vision (the PRL) in individuals with central vision leads to more idiosyncratic functional connectivity maps the part of cortex associated with that part of vision. ***p<0.001 for ANOVA of group by ROI.

Next, we compared similarity scores from only the cPRL region, using an independent samples t-test. Our results showed that the similarity of cPRL-seeded functional connections was higher in healthy controls than in MD participants (Figure 5, t(5794)=14.8328, p<0.001). These results indicate that experience with increased usage is associated with more idiosyncratic functional connection patterns.

**Figure 5.**
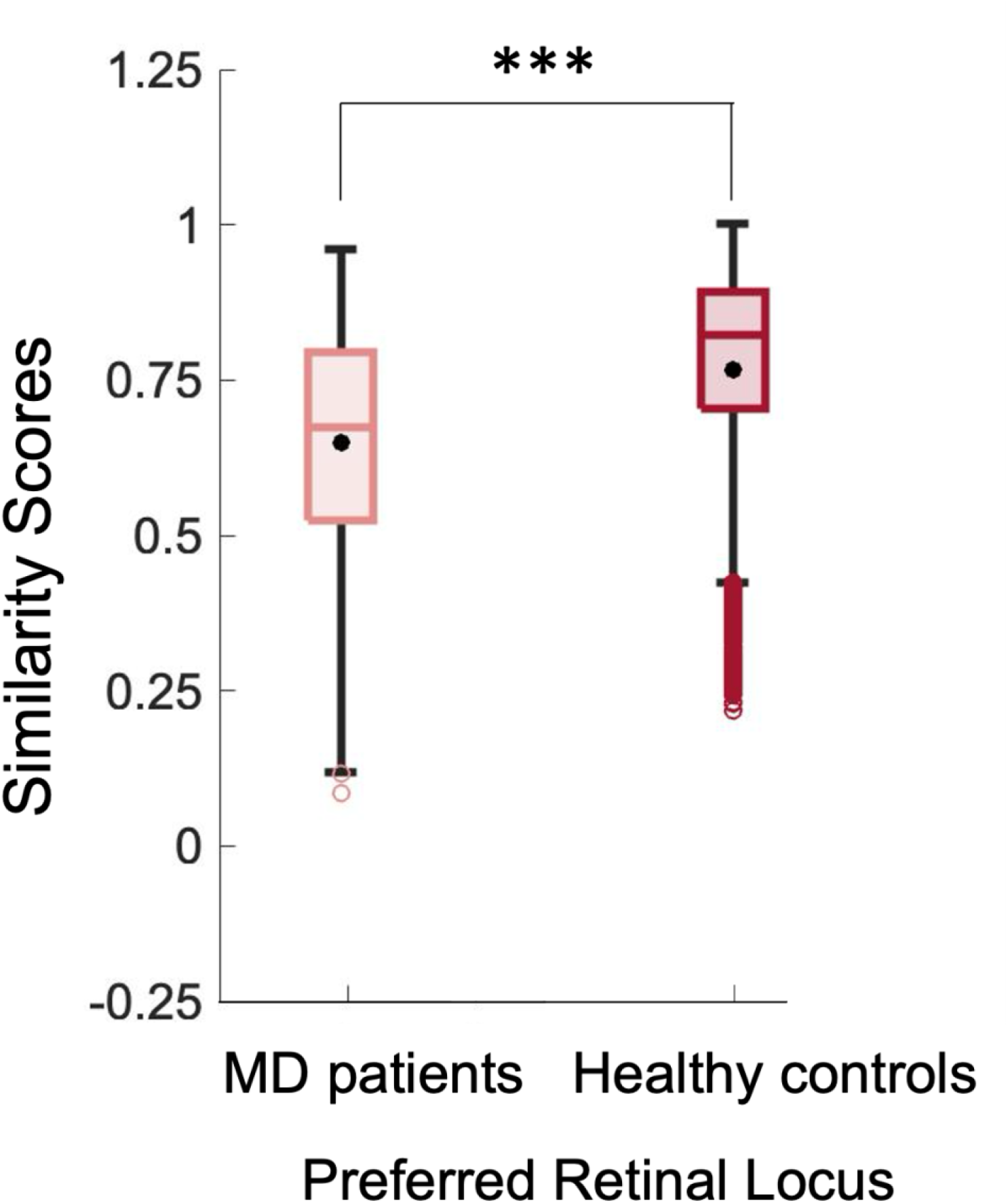
Cortical projection of the cPRL-seeded FC maps is more idiosyncratic in MD patients than in the healthy vision control group. Group-level independent sample t-test comparisons of similarity scores showed that cPRL (t(5794)=14.8328, p<0.001) seeded FC maps are have lower similarity scores (more idiosyncratic) for MD patients than healthy vision controls. Colored data points indicate individual measurements. The black data point represents the group average, and the colored horizontal line represents the group median. Error bars show ±1 standard error. Top and bottom bounds of each box represent the 75th and 25th percentiles, respectively. The whiskers extend to the minima and maxima data points, not considered outliers. ***p<0.001.

In order to identify whether the distributions of FC maps for the PRL were becoming more like those of central vision, we analyzed individual variability in LPZ and cPRL-seeded FC patterns. Figure 6 compares the distributions of FC map similarities for LPZ and PRL for both groups. These are the same values as shown in 3 and 4, just directly compared here. Interaction between ROI and participant diagnosis was statistically significant (F(1,6528)=267.34, p<0.001). Our results indicate that increased use of peripheral vision in people with central vision loss makes their patterns of functional connections more idiosyncratic, and more comparable to the idiosyncratic functional connections that are typical for central vision. These findings are consistent with the idea that as we use specific functional connections more, they become more different in their own way.

**Figure 6.**
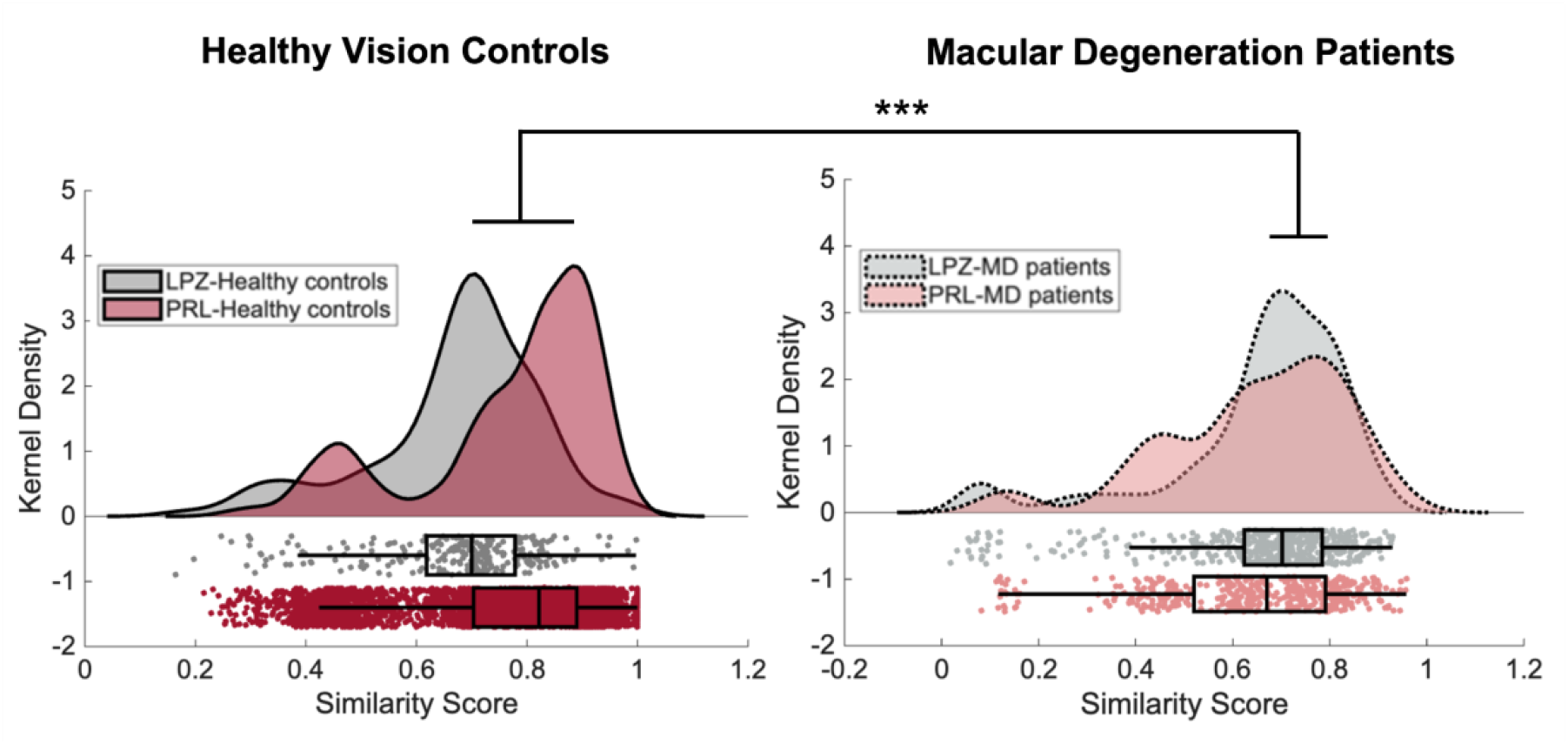
In MD patients, the cPRL has a pattern of idiosyncrasy like that in central vision. LPZ and cPRL-seeded FC maps’ similarity score distributions are significantly different between healthy vision controls and macular degeneration patients. Interaction between ROI and participant diagnosis is statistically significant (F(1,6528)=267.34 p<0.001). LPZ (shown in gray) and cPRL (shown in red) -seeded FC maps’ distribution of similarity scores are shown for macular degeneration patients (dashed line) and healthy vision controls (solid line). ***p<0.001 for ANOVA of group by ROI.

### Standard Group Comparisons Showed Limited Evidence For Plasticity After Central Vision Loss

Standard group-level comparisons can reveal differences between groups; however, individual variability is high in the MD patient population. We analyzed standard group-level differences between the MD and healthy control groups to test outcomes when we ignore individual variability. Independent sample t statistics for LPZ and cURL regions are shown in Figure 7. Our group-level LPZ, cPRL, and cURL-seeded FC map comparisons revealed that only the LPZ FC map is different between the groups; healthy vision controls have stronger functional connectivity to a cluster in the temporo-parieto-occipital cortex compared to MD patients (Figure 7B, cluster size=6031, FWE-corrected p=0.022). Figure 7B shows that MD patients who have central vision loss have weaker functional connections from the central vision processing region to regions within temporal, parietal and occipital lobes. This is consistent with previous findings (17, 18) and suggests that loss of use of a brain area due to sensory deprivation results in connectivity decreases that are consistent across people. However, using this standard method, we could not detect any group difference in functional connections of more attentive usage of a specific area in periphery (PRL) between MD patients and healthy vision controls (Figure 7C & D). Thus, the standard group comparison method showed limited evidence for plasticity of functional connections to the deprived brain area, and no evidence for plasticity of functional connections to the ‘more attended’ brain area.

**Figure 7.**
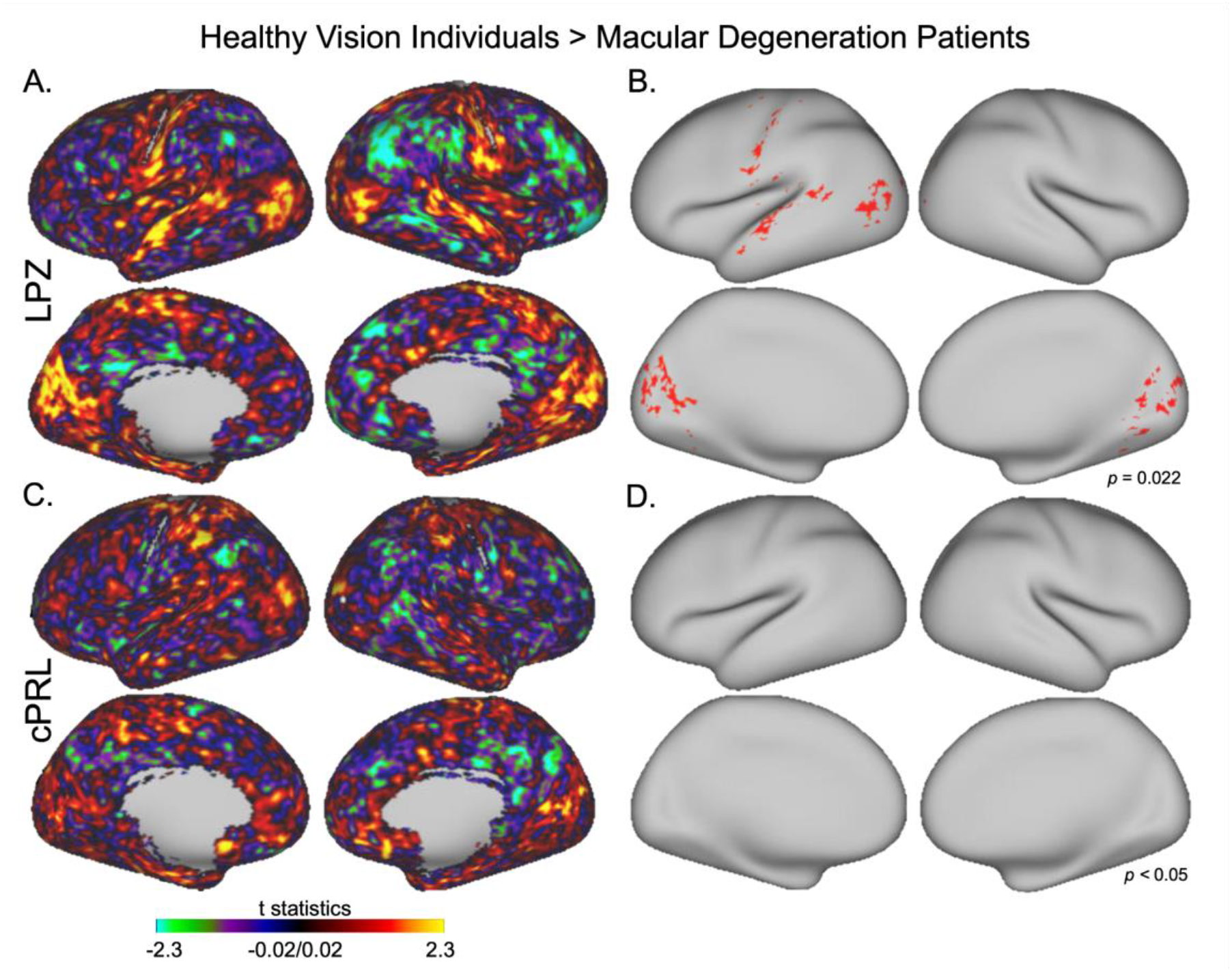
Group-wise comparison of whole-brain FC maps for Macular Degeneration patients vs. healthy vision control groups. Group-level comparisons showed limited evidence for plasticity, when measured as a mean difference between groups. A & C. Independent-sample t-test scores between healthy vision individuals and MD patients for whole brain FC maps for LPZ (A) and cPRL (C), Brighter colors represent more significant results. B & D. FWE-corrected p values between healthy vision individuals and MD patients for LPZ (B) and cPRL (D). (B) shows that the healthy vision control group showed stronger functional connectivity in HC than MD participants to a cluster including temporo-parieto-occipital regions (cluster size=6031, FWE-corrected p=0.022). Group level analysis results were projected onto to 32k cifti space template. Note that there are no significant voxels in D, indicating no significant between-group differences in connection to the cortical projection of the PRL, when measured this way.

## Discussion

In this study, we examined the idiosyncrasy of whole brain functional connections in two models; healthy individuals – who preferentially attend to their central vision more than their peripheral vision – and patients with central vision loss – who preferentially attend to a specific peripheral vision area more than other areas of vision. In participants with healthy vision, we found that central vision representations had much more idiosyncratic cortical connection patterns than peripheral vision. In participants with central vision loss, we found that the representations of the preferred peripheral retinal locations (PRL) had much more idiosyncratic cortical connection patterns than other parts of peripheral vision. Together, these results strongly suggest that cortical regions that experience frequent attentive use develop idiosyncratic cortical connections.

### Central vision FC maps are more idiosyncratic than peripheral vision FC maps

Central and peripheral vision have different functional roles. High acuity visual information from the central few degrees of the visual field is critical for everyday activities such as reading and object identification (19–21), whereas peripheral vision is essential for visual tasks such as visual search and getting the gist of the scene (19, 22). These functional differences between central and peripheral vision are essential for typical use. For example, peripheral visual information is used to plan saccadic eye movements which orient high-acuity central vision and attention to a new location. Thus, we may expect that the patterns of brain connections for the representations of central vision would differ from the patterns of connections for central vision, despite the fact that those representations may reside within the same cortical area (for example, V1). In fact, although the bulk of connections across V1 are relatively similar, connections between high-order brain areas and lower-level visual processing areas (e.g. the primary visual cortex, V1) are more robust in the central than peripheral visual fields (23–25). If cortical feedback resources are more available to the central visual field, processing of information from the peripheral visual field might suffer from insufficient feedback for input verification or disambiguation. This suggests that peripheral vision would be vulnerable to illusions associated with lack of verification from top-down signals, and indeed some illusions follow this pattern and are present in peripheral viewing but not central viewing (23, 26–28).

While the connection patterns do differ, on average, between central and peripheral vision representations, as can be seen in previous studies (24, 25), the effect sizes in previous studies are relatively modest. On the other hand, when we examine the whole-brain patterns, we see that central representations show extremely idiosyncratic connection patterns, relative to peripheral representations. Interestingly, previous studies that have examined cross-subject variability in connection patterns have found that the occipital lobe in general has relatively little inter-subject variability (5). Cortical networks associated with attention and executive function, such as the Fronto-parietal network, and the ventral attention network have the most idiosyncratic patterns of connections, while visual cortical and motor cortical networks have the least idiosyncratic patterns overall (e.g., Figure 2 in reference (5)). This suggests that brain regions involved in attention and executive functioning develop idiosyncratic patterns of connections. This is consistent with our finding that central vision, which is used daily for tasks involving attention, has more idiosyncratic connection patterns. Our finding that the PRL showed more idiosyncratic connections after central vision loss suggests that some degree of this region-to-region difference in connection patterns is due to experience, and is not solely driven by evolutionary pressures.

### Cortical regions associated with increased use have more idiosyncratic FC maps

Many people with central vision loss preferentially use a specific part of their intact retina, the preferred retinal locus (PRL), for daily tasks. Our results suggest that when people with central vision loss use this PRL for attention-demanding tasks, the patterns of connections become more idiosyncratic, less stereotypical. This result is consistent with the general concept that more attentive use leads to more idiosyncratic connection patterns.

Previous studies using standard analytical approaches revealed that patients with central vision loss show enhanced functional connectivity between peripheral early visual cortex (representing the PRL) and higher-order regions involved in attention and oculomotor control, including the intraparietal sulcus and frontal eye fields (17, 29). These reorganized connectivity patterns echo the cortico-cortical interactions observed during central vision processing in normally sighted individuals, which are characterized by strong coupling between foveal V1 and attentional networks (7, 30). The similarity in network-level profiles suggests that when peripheral vision becomes the dominant source of visual input, the brain adapts by allocating cognitive and attentional resources in a way that functionally substitutes for the lost foveal input. In addition, Mueller et al (2013) argue that higher cross-subject variability (more idiosyncratic connections) is associated with stronger long-distance connections. This is intriguing since previous data show that central representations in V1 have stronger long-distance connections to lateral frontal regions than peripheral representations (25). These results suggest the possibility that these frontal connections are the source of the cross-subject differences. This alteration in functional connections likely reflects both experience-dependent plasticity and task-driven recruitment, highlighting the adaptability of visual-attentional systems to changing sensory demands.

### Idiosyncratic Reorganization Reflects Use-Dependent Plasticity in Adulthood

Our findings provide new evidence that functional connections of V1 retain the capacity for experience-dependent plasticity well into adulthood. The capacity for plasticity of V1 in adulthood has been the subject of some controversy (31–33). A great deal of literature in human participants has suggested that the primary visual cortex has extremely limited plasticity in adulthood (18, 31, 32, 34, 35). For example, studies that examine whether there is remapping of visual representations to new locations have shown that there is very limited if any shifts of representations (32, but see 33, 35–37), though some studies in animal models show that the receptive field centers do shift to some degree (31, 38). Studies that examine functional connections to V1 after loss of vision have shown preserved patterns of local connections in patients with vision loss (34, 39). Studies of more distant functional connection patterns have shown limited shifts in connection patterns in adulthood (17, 18). Data examining mean differences between groups in Figure 7 are consistent with these general findings; we saw limited loss of connections to the LPZ (a cortical region that had lost sensory inputs), and no significant changes in connections to the PRL (a cortical region that showed increased attentive use) after multiple comparisons correction. These studies show that it is difficult to observe evidence of plasticity in visual cortex in adulthood using standard approaches. However, learning does occur in adulthood, and it stands to reason that we may be able to observe plasticity associated with that learning. By examining whole brain functional connection patterns by taking individual differences into account, using the idiosyncrasy approach we use here (Figure 1), we can observe strong evidence of plastic changes that are different person-to-person (Figure 6). This approach shows that functional connections of V1 are plastic in adulthood, and specifically, that increased attentive usage drives that plasticity. These results also suggest that one potential aspect contributing to the controversy regarding plasticity is that most studies in humans have compared means of one group to another, rather than taking individual differences into account.

It is worth noting that a large number of studies have examined adult plasticity of many aspects of neural activity and connections. While several studies suggest there is limited remapping of the centers of receptive fields after vision loss (31, 32, 35, 36), there is strong evidence that portions of the LPZ shows BOLD activity during performance of attention-demanding tasks (37, 40, 41). This activity, however, is not present during passive fixation (37, 42), is not specific to any given spared portion of retina (like the PRL) (41), and is present during non-visual tasks (42). Further, studies of the brains of people who are congenitally blind have shown some brain connectivity patterns that are consistent with the literature here (43–46). These results suggest that the LPZ activity in patients with central vision loss results from something other than direct input from the thalamus, and may not reflect retinotopic sensory processing. This suggests that there is some degree of plasticity of possibly lateral or top-down connections to LPZ, consistent with some models (47). Less work has examined the neural responses of brain areas associated with increases in attentive vision, such as the PRL. Some studies have suggested that there is an increase in cortical thickness in the cortical representation of the PRL (48–50). Furthermore, a previous study from our team (18) directly examined group-wise differences in connections to that area, and found considerable subject-to-subject variability in the strength of connections in the patient group, a result which inspired the current study.

### Sensory deprivation is associated with no change in idiosyncrasy

Whereas our data show strong evidence that connection patterns become more idiosyncratic with attentive use, we do not see evidence of decreases in idiosyncrasy following lack of use. Figure 3C shows that the functional connection patterns for the LPZ for those who have lost sensory input to the LPZ are similarly idiosyncratic to healthy vision controls. Thus there is no change in idiosyncrasy of connections associated with decreased attentive use of the region. This is surprising, given that by far the biggest visual experience change that patients had was a loss of inputs to this central vision region. This suggests that decreased attentive use does not alter the idiosyncrasy of connections.

Interestingly, MD patients did show modest declines relative to the mean of healthy vision controls in strength of connections between the LPZ and several brain areas shown in Figure 7B. Conversely, strengths of connections to the cPRL did not significantly differ. This pattern of results suggests that while increased attentive use may lead to more idiosyncratic patterns of connections, decreased use leads instead to weakening connections. This weakening happens in a way that does not change the idiosyncrasy of the whole-brain patterns: they are just as variable, just weaker. These results suggest that the network mechanisms that result in plasticity after increased attentive use and decreased attentive use are distinct. The results are consistent with the idea that increased attentive use strengthens some connections and weakens others, as needed for the specific and idiosyncratic functions needed for the attended task. On the other hand, decreases in attentive use (as in the LPZ) may lead to only weakening of connections, and no appreciable change in the spatial pattern of connections, relative to a typical brain.

To sum up, our results showed that experience with gain of attentive use in the periphery leads to more idiosyncrasy, a.k.a. more variability between MD patients and healthy vision controls, and this effect is specific to cPRL connections.

### Individual-specific approaches are necessary for disease populations

Generally, neuroimaging studies aggregate small quantities of functional data (5-10 min) across many individuals to characterize brain function and connectivity. Due to the low temporal signal-to-noise ratio of functional data (51), the general tendency in the neuroimaging literature is to identify group-level central tendencies that generalize across individuals to overcome small quantities of functional data from individuals, but this approach ignores subject-specific features. The necessity of implementing individual-specific approaches rather than relying solely on group comparisons becomes particularly evident in populations with high inter-individual variability, like many patient populations, including those with central vision loss. The impact of central vision loss on the brain’s functional architecture can vary significantly between individuals(52). By acquiring relatively high amounts of functional data per participant (four runs), we were able to implement individual-specific approaches that allow for a more nuanced understanding of the neural adaptations occurring in response to central vision loss. Tailoring interventions and rehabilitation strategies to each individual’s unique neuroplastic profile becomes a possibility using these types of approaches, and standardized group-based approaches may not fully capture the diversity of neural responses and compensatory mechanisms at play in the context of visual impairment. This individualized perspective is crucial for advancing personalized and effective interventions and rehabilitation strategies for those with central vision loss.

### Ruling out potential confounding factors that might lead to inter-individual variability in functional connections

Resting-state data are by nature heterogeneous (5, 53). It is therefore essential to rule out confounding factors that might lead to incorrect interpretations of our measures of idiosyncrasy. In this study, we controlled some conditions that might affect the resting state blood oxygen level dependent data during the acquisition to avoid confounds. Conditions such as participants closing their eyes vs. having their eyes open, or fixating vs. moving their eyes influence intrinsic connectivity in resting state data (54). We asked all of our participants to keep their eyes open, fixate on the cross at the center, and let their minds wander throughout the scans. We monitored their eye position and eye opening using an infrared illuminator and camera during data acquisition in order to assess subjects’ compliance. Other factors may also contribute to the variability of functional connectivity patterns. Specifically, factors like performance of a task just prior (55), caffeine intake (56, 57), and falling asleep (58) can bias resting state measurements. Also, since Macular Degeneration is a retinal disease that limits light input to the retina, the existence of light in the scanner room might be a confounding variable across the MD participants and between the groups. In our study, we collected our resting-state functional data in complete darkness to eliminate any potential differences in visual sensory input across participants. Moreover, we also kept the order of the data acquisition the same for all participants, controlling for the prior task, and as described above, tracked the participants’ eyes during the scan. We controlled these possible confounds as much as possible to reduce variance between the participants that was evoked by factors other than intrinsic connectivity differences between the groups.

### Across-subject variability of the location of the PRL

Each MD patient develops a PRL at a different retinal location, due to many factors including the extent and shape of their retinal scotoma. Patients do not always choose the optimal PRL location for the best visual acuity (59). MD patients often develop PRLs at a retinal location considered “unfavorable for reading”, usually to the left of the retinal lesion, corresponding to the left visual field of the central vision loss (60, 61). Fixation to the left side of the scotoma might be related to left-to-right reading in the Western World, this is also consistent with the findings in the Arab population, who read from right-to-left, and preferentially and have PRLs to the right of the scotoma (61). Visual rehabilitation studies have shown that when individuals were trained above or below the retinal lesion, their performance improved (62, 63). When we checked PRL locations in our patient population, MD patients developed PRL in a variety of retinal areas that correspond to mostly lower and left visual fields (see Supplementary Materials 2). In the literature, it has been shown that perceptual performance varies across different visual field locations (64, 65). Studies showed that horizontal visual field asymmetries may reflect functional hemispheric differences (66). There are also reliable behavior/performance differences between the upper and lower visual fields (67–69). The lower visual field is advantageous in target detection and discrimination (64, 65, 69), visual acuity (70), contrast sensitivity (71), spatial attention (72), and perception of illusory contours (73). Because of the different personal experiences and needs for daily tasks, usage preferences of the specific intact areas in the periphery might differ from MD patient to MD patient. One of the benefits of our analytical approach is that it takes these different PRL locations into consideration and allows us to examine connectivity patterns to a specific swath of cortex corresponding to an individual patient’s PRL.

## Materials and Methods

### Participants

MRI data were acquired from 44 (21 MD, 23 control) participants as a part of the Connectomes in Human Diseases Macular Degeneration Plasticity project (74). MD participants had been previously diagnosed by an ophthalmologist as having bilateral macular degeneration. Our inclusion criteria for the MD group include having a diagnosis for at least 2 years prior to being enrolled in the study, having bilateral dense scotomas at the center of the fovea, and not having any psychiatric or neurological diseases. All MD participants underwent visual acuity testing, using the Early Treatment Diabetic Retinopathy Study (ETDRS) test (75), macular integrity assessment (MAIA, Centervue, Padova, Italy), and Optical Coherence Tomography (OCT) to confirm that all MD participants have foveal scotomas in both eyes and low visual acuity. Demographic information of MD participants is summarized in Table 1. Written informed consent was obtained from all participants prior to enrollment in the study. Data were collected in accordance with the ethical standards under the oversight of the University of Alabama at Birmingham (UAB) Institutional Review Board.

**Table 1.**
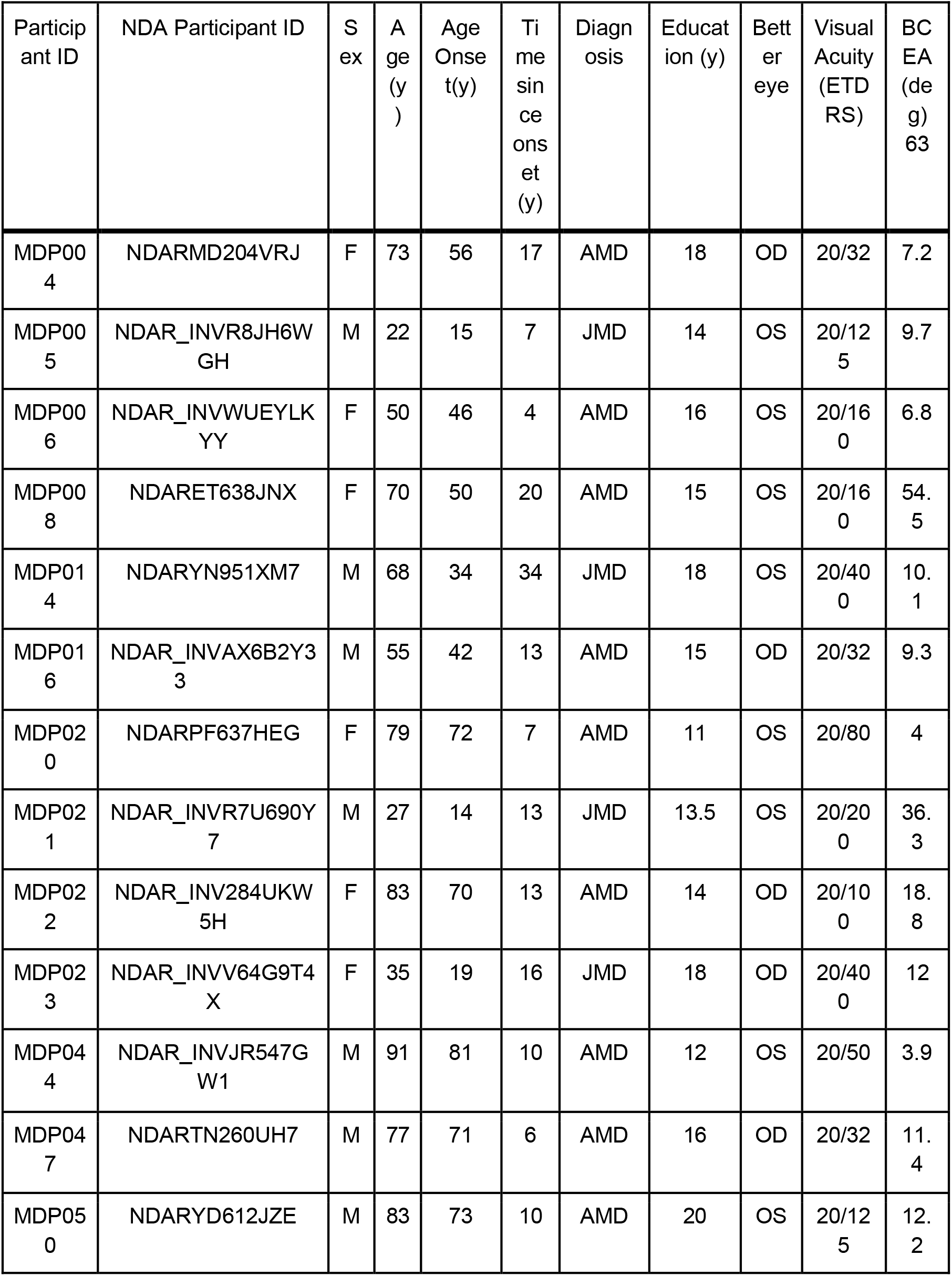
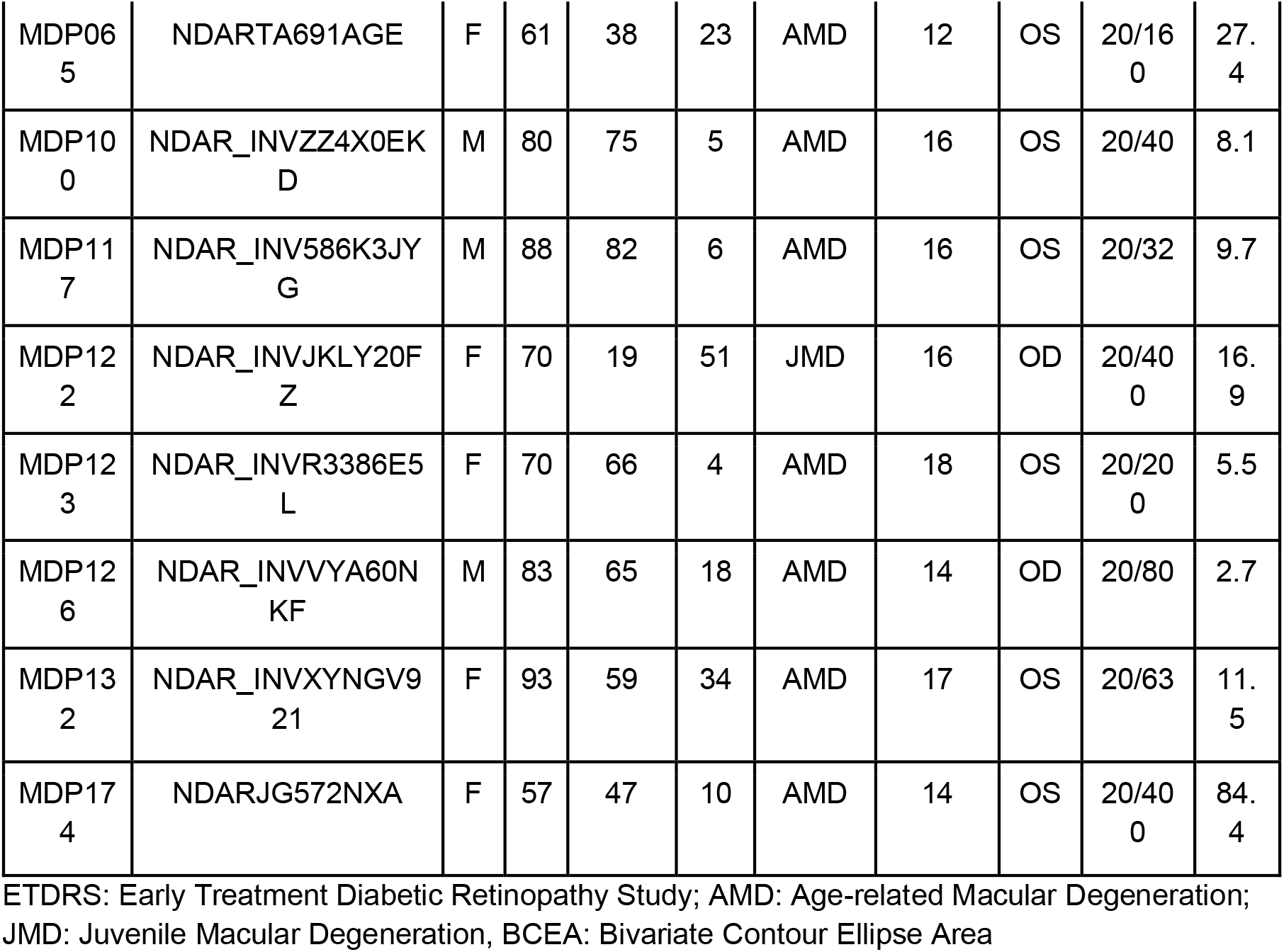
Demographic information of participants with Macular Degeneration.

### 2.2 MRI Data Acquisition

All images were acquired on a 3T Siemens Prisma scanner using a 64-channel head-coil at the University of Alabama at Birmingham, Alabama using protocols based on those of the Human Connectome Project (74, 76). High-resolution 3D anatomical scans were obtained (T1-weighted; repetition time (TR) = 2400 ms; echo time (TE) = 2.22 ms; field of view [FOV(ap,rl,fh)] = 208 x 208 x 144 mm; voxel size = 0.8 mm isotropic; flip angle (FA) = 8 deg). T1w images were reacquired if the initial acquisition was of poor quality. Resting-state functional scans (eyes open) were acquired in total darkness (T2*-weighted, TR=800 ms, TE=37 ms, FA=52º; voxel size=2 mm isotropic; echo spacing=0.58 ms; multi-band acceleration factor = 8), resulting in 420 volumes per scan. To ensure complete darkness, the investigators blocked out all possible sources of light in the room (windows, waveguides, lights on equipment) prior to the start of the resting-state scan. Participants were instructed to relax and keep their eyes open until the scan was complete. An infrared eye-tracking system camera (M0203 Livetrack AV, Cambridge Research Systems), which does not emit visible light, was used to monitor participants during scanning to ensure that their eyes remained open during the scanning session. The resting-state data were acquired in four runs and each run took 5 min 36 sec.

### 2.3 MRI Data Preprocessing

Raw data fMRI data files were formatted according to the Brain Imaging Data Structure (BIDS) standard (77) to enable use with open-source pre-processing pipelines. Initial first-pass quality control was performed through manual inspection of the data using MRIQC (78). Preprocessing of the fMRI data was performed using fMRIPrep 1.2.5 (79), which is based on Nipype 1.1.6 (80).

#### 2.3.1 Anatomical data preprocessing

T1-weighted images were corrected for intensity non-uniformity and skull-stripped. Brain surface reconstruction was performed using recon-all from FreeSurfer v6.0.0 (81). Spatial normalization to the ICBM 152 Nonlinear Asymmetrical template version 2009c was performed through nonlinear registration with antsRegistration (ANTs 2.2.0, (82)), using brain-extracted versions of both T1w volume and template. Brain tissue segmentation of cerebrospinal fluid (CSF), white matter (WM), and gray matter (GM) was performed on the brain-extracted T1w using FSL 5.0.9 (83).

#### 2.3.2 Functional data preprocessing

Functional MRI data preprocessing in fMRIPrep was carried out as follows: First, a reference volume and its skull-stripped version were generated using fMRIPrep. The BOLD reference was then co-registered to the T1w reference using bbregister (FreeSurfer), which implements boundary-based registration (84). Co-registration was configured with nine degrees of freedom to account for distortions remaining in the BOLD reference. Head-motion parameters concerning the BOLD reference (transformation matrices and six corresponding rotation and translation parameters) are estimated before any spatiotemporal filtering (85). BOLD runs were slice-time corrected using 3dTshift from AFNI 20160207 (86). The slice-time corrected BOLD time-series were resampled onto their original, native space by applying a single composite transform to correct for head motion and susceptibility distortions. These resampled BOLD time series will be referred to as preprocessed BOLD in original space, or just preprocessed BOLD. The BOLD time series were also resampled to MNI152NLin2009cAsym standard space. First, a reference volume and its skull-stripped version were generated using fMRIPrep. Several confounding time series were calculated based on the preprocessed BOLD: framewise displacement (FD), DVARS, and three region-wise global signals. FD and DVARS were calculated for each functional run, using their implementations in *Nipype* 1.1.6 (80) following the definitions by (87). The three global signals are extracted from the CSF, the WM, and the whole-brain masks.

Additionally, a set of physiological regressors was extracted to allow for component-based noise correction (88). Principal components are estimated after high-pass filtering the preprocessed BOLD time series (using a discrete cosine filter with 128s cut-off) for the two CompCor variants: temporal (tCompCor) and anatomical (aCompCor). Six tCompCor components are then calculated from the top 5% variable voxels within a mask covering the subcortical regions. This subcortical mask is obtained by heavily eroding the brain mask, which ensures it does not include cortical GM regions. For aCompCor, six components are calculated within the intersection of the aforementioned mask, and the union of CSF and WM masks is calculated in T1w space after their projection to the native space of each functional run (using the inverse BOLD-to-T1w transformation). The head-motion estimates calculated in the correction step were also placed within the corresponding confounds file. All resamplings were performed with a single interpolation step by composing all the pertinent transformations (i.e. head-motion transform matrices, susceptibility distortion correction when available, and co-registrations to anatomical and template spaces). Gridded (volumetric) resamplings were performed using antsApplyTransforms (ANTs), configured with Lanczos interpolation to minimize the smoothing effects of other kernels (89). Non-gridded (surface) resamplings were performed using mri_vol2surf (FreeSurfer).

#### 2.3.3 Motion Correction

Motion artifacts during fMRI scanning are known to produce substantial effects on functional connectivity data (90, 91). To mitigate such confounds, we performed motion scrubbing on the preprocessed data from fMRIPrep using the XCP engine workflow (92) using a framewise displacement threshold of 0.5mm. Following the preprocessing procedure, the data were converted into a template space (fsLR_32k cifti-space) using a combination of Ciftify (93) and the Connectome Workbench (94).

### 2.4 Region of Interest Definitions

Regions of interest in the primary visual cortex (V1) were defined based on standard atlases of visual regions (95) coupled with maps of the visual field that were acquired from retinal imaging as described in our previous work (96) (Figure 8). An LPZ region of interest was created for each participant that included the 100 vertices on each hemisphere with the lowest eccentricity values (200 total vertices per participant). We estimated the location at which the MD patients tended to primarily fixate (i.e., - the preferred retinal locus, PRL) from macular integrity assessment (MAIA). The PRL (Figure 8A) was defined based on the cloud of fixation locations over time (Figure 8B). The center of the PRL was defined as the center of the cloud, and the boundary was defined as the Bivariate contour ellipse area that included 63% of fixation locations (97). We also defined a control region that we refer to here as the cortical representation of the “Un-preferred Retinal Locus” or URL. This region was defined by identifying a region on the retina outside of the lesion with the same eccentricity as the PRL region. We used MD participant-specific cPRL and cURL region region of interests for all healthy vision participants. Foveal (scotoma) and peripheral retinal locations (PRL and URL) for each MD participant were identified from retinal images, and they were mapped on the cortex by using the cortical mapping method (Figure 8 C&D). Details of the cortical mapping method were described in Defenderfer et al, 2021 (96). In brief, we created binary LPZ, PRL and URL ROIs in retinal space using Photoshop.Standard retinotopic atlases including eccentricity, polar angle and sigma were transferred to each participant’s cortical surfaces through the Python library neuropythy (95). These atlases assign visual eccentricity, polar angle and sigma values to each vertex in early visual areas. Each pixel in the LPZ, PRL and URL images was assigned an eccentricity and polar angle value based on pixel distance from the center of the fovea ROI multiplied by a pixel to visual degree conversion factor obtained from the MAIA microperimeter. In our study, for each vertex in V1, the retinotopic coordinates were extracted from the retinotopic atlases, and the pixel with the closest eccentricity and polar angle values was found. If that pixel was located in the ant given ROI, that vertex was added to that specific ROI label. All ROIs were mapped on individuals’ native cifti space and then converted into 32k standard space.

**Figure 8.**
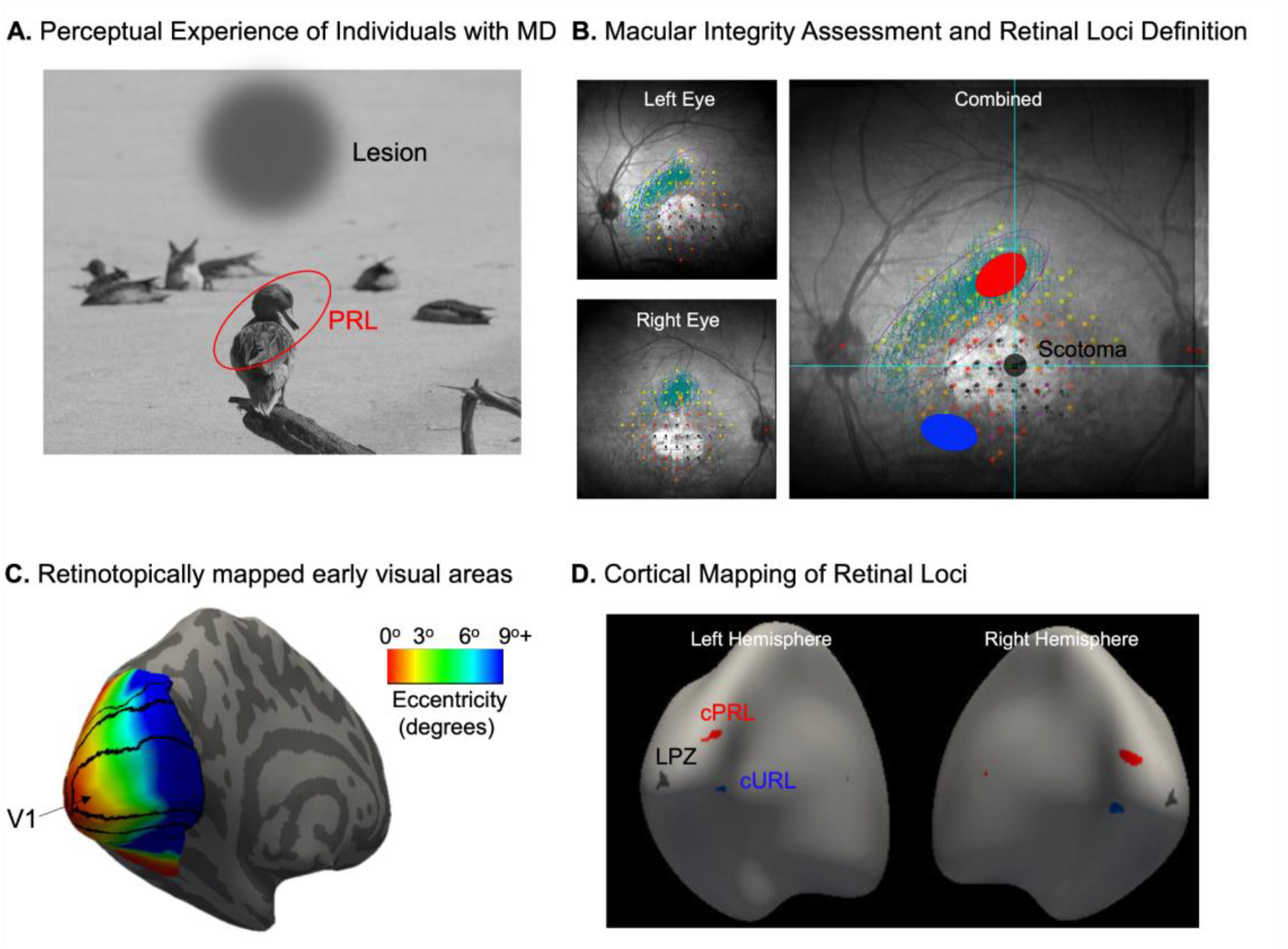
Identifying retinal loci (Lesion, PRL, and URL) and their cortical projections. (A) Representation of the perceptual experience of individuals with Macular Degeneration (MD). (B) Fundus photograph of the retina from one eye of a representative participant with central vision loss. The grayscale background shows the anatomy of the retina. Microperimetry results are shown in the grid of colored dots. The colors of dots represent the threshold for detection of a spot of light at that location on the retina. Black dots indicate no light detection. Teal dots represent retinal locations to which the participant orients a fixation target. Cyan ellipse encloses 63% of participants’ fixations. Note that the right eye has a tighter fixation window. The location of these fixations on the better eye is defined as the Preferred Retinal Location (PRL). The Unpreferred Retinal Location (URL) is defined in an equally eccentric location to the PRL, with similar visual detection and retinal health. (C) Illustration of the anatomic atlases of eccentricity, polar angle, and sigma mapping in early visual areas. Eccentricity is represented across the cortex with central vision at the occipital pole. (D) Based on the anatomic atlases, we define the cortical projections of the scotoma (LPZ, shown in dark gray), cPRL (shown in red), and cURL(shown in blue) in V1. The LPZ region of interest was defined as the 100 vertices with the lowest assigned eccentricity values in both hemispheres in V1.

### 2.5 Functional Connectivity Analysis

The average time course of each ROI (LPZ, cPRL and cURL) was extracted from the preprocessed resting state fMRI data using in-house MATLAB scripts. Functional connectivity was then calculated as Pearson’s correlation between the timecourses of each ROI (seed) to each voxel in the brain for all individuals. The resulting whole-brain seed-based correlation maps are referred to here as “FC maps”. Then, we performed Pearson’s correlation comparison between healthy vision participants and other healthy vision participants to estimate the ‘idiosyncrasy’ of the functional connectivity patterns in healthy individuals. We also performed Pearson’s correlation analysis between MD patients and healthy vision participants to understand the similarity in functional connectivity FC maps between groups. Next, we compared the similarity of the MD patients vs healthy vision participants and healthy vision participants vs. healthy vision participants to discuss altered individual experience effects on typicalities of the functional connectivities based on LPZ, cPRL, and cURL areas. In these analyses, all Pearson’s correlation results were converted to Fisher’s z scores.

### 2.6 Statistical Analysis

We followed both individual-specific (Figure 1) and group-level (Figure 9) approaches to evaluate experience-dependent plasticity in whole-brain functional connectivity in individuals with MD.

**Figure 9.**
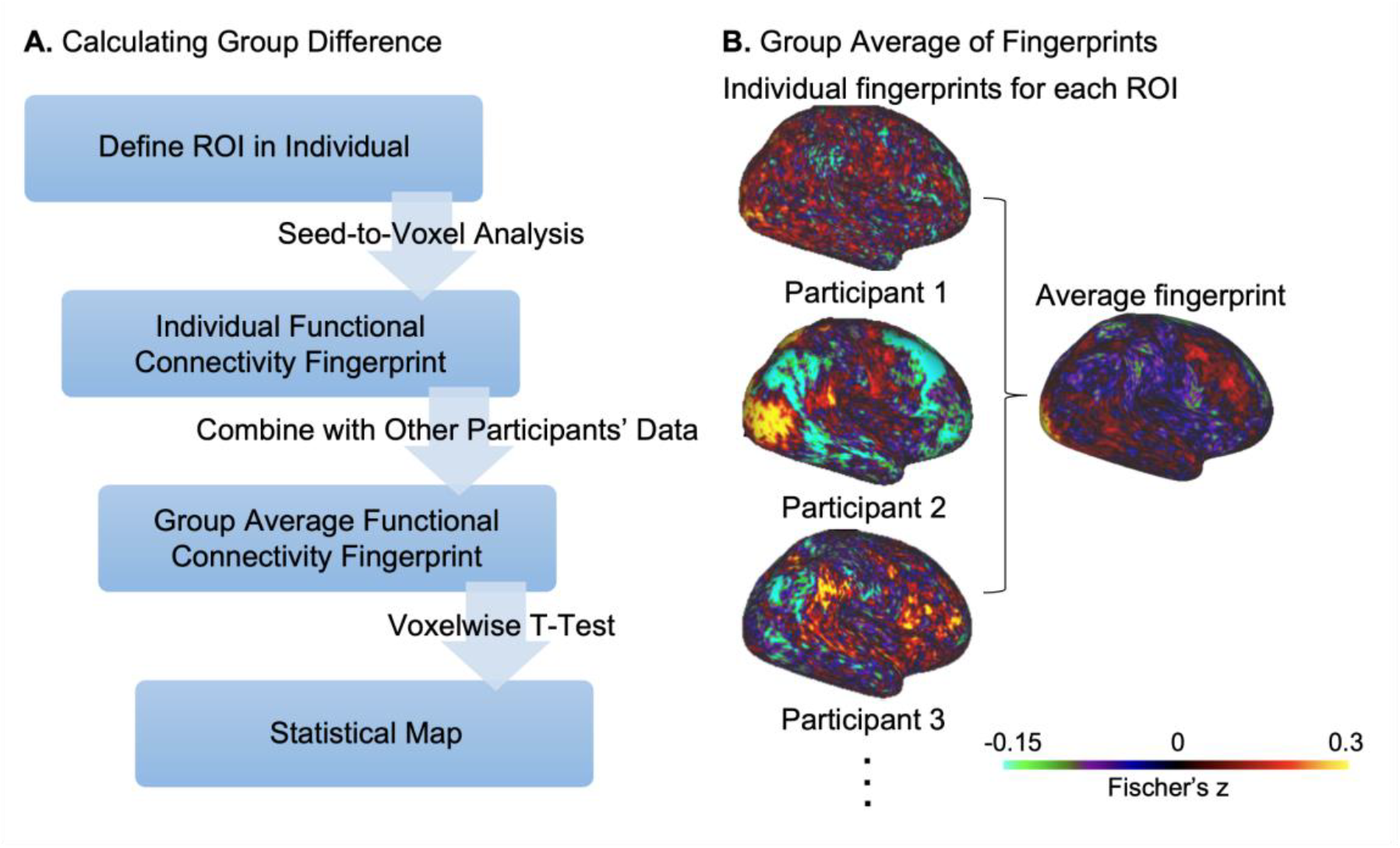
Schematic representation of (A) group comparison analysis flow and (B) group averages seed-to-voxel functional connectivity map analysis. Functional connectivity group averages were calculated for both MD patients and healthy vision participants for LPZ, cPRL, and cURL regions of interest. LPZ-seeded FC maps were defined as the 100 vertices with the lowest assigned eccentricity values around the fovea in both hemispheres in V1. PRL and URL-seeded FC maps were calculated based on each MD participant’s specific regions of interest.

#### 2.6.1 Defining Idiosyncrasy of a Functional Connectivity (FC) Map

We defined scotoma, PRL and URL ROIs in individuals’ MAIA images then we mapped them on their cortices. By using LPZ, cPRL, and cURL as seeds we performed seed-to-voxel analysis and created individual FC maps for each individual. Next, we compared each FC map to FC maps of each healthy control by performing Pearson’s correlation. Then, we graphed histograms of similarity scores for each group (Figure 1).

For the individual-level comparisons, we performed similarity analysis by performing Pearson’s correlation analysis between each healthy vision control and every other healthy vision control to estimate typical functional brain connectivity patterns. We performed an analysis of the variance (ANOVA) of linear mixed-effects models to compare inter-individual variability with factors of ROI (LPZ, cPRL, and cURL) for within-subject variability comparisons in healthy vision controls. We extracted similarity values and fit them in our mixed-effect model. This model included an ROI fixed effect predictor and a random effect for participants, as well as a random effect for the MD patient from whom the ROI was defined. This model is summarized in the following formula:

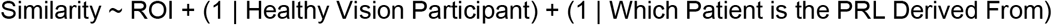

We performed an analysis of the variance (ANOVA) of the linear mixed effect model with factors of ROI (LPZ, cPRL, and cURL) and diagnosis (MD vs. healthy vision participants) to compare inter-individual variability between the groups. We derived similarity values and fit them in our mixed effect model. We analyzed main effects of ROI and diagnosis of the participant, and interaction between diagnosis of participant and ROI as fixed effect predictors and random effects for Participant, and MD participants that we defined ROIs from by using the following formula:

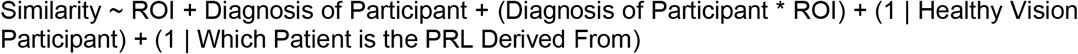

#### 2.6.2 Group-level comparisons

We also completed a more standard group-level analysis (Figure 9) to compare functional connectivity FC maps in individuals with MD to controls. For each MD participant, the PRL and URL were defined based on their own retinal data, as described above. For each healthy vision control participant, the cPRL from every MD participant was defined on that control participant’s cortex, and an FC map was made for each of those ROIs. Thus, each control participant contributed multiple FC maps to the analysis, one for each cPRL. The same is true for the cURL areas. We used an independent sample t-test to compare the FC maps for ROIs between MD patients and healthy vision participants. Since our MD participants have bilateral scotomas of different sizes, we identified and compared FC maps from 100 vertices of the cortical region corresponding to central vision in each hemisphere to keep the size of the LPZ region consistent across individuals. We used a cluster-based thresholding method for group-level comparisons (98, 99). We corrected our results for multiple comparisons and we set the threshold to 1.5 for t contrasts. We performed group-level analyses for ROIs from PRL, URL and LPZ.

## Acknowledgments

We would like to thank the Connectomes in Human Diseases Grant to KMV NIH NEI 1 U01 EY025858-01A1, and R01EY031589, UAB Center for Clinical and Translational Science UL1 TR000165, Vision Science Research Center P30 EY003039, Civitan International Research Center, McKnight Brain Research Foundation, Edward R. Roybal Center for Translational Research on Aging and Mobility NIA 2, P30 AG022838, UAB Comprehensive Center for Healthy Aging, for providing support for this study.

## References

1. S. Marek, et al., Reproducible brain-wide association studies require thousands of individuals. Nature 603, 654–660 (2022).

2. G. MacDonald, Great Expectations, Episode 2505, Segment 2. 2505: Great Expectations (2025) (August 27, 2025).

3. J. S. Damoiseaux, et al., Consistent resting-state networks across healthy subjects. Proc Natl Acad Sci USA 103, 13848–13853 (2006).

4. J. D. Power, et al., Functional network organization of the human brain. Neuron 72, 665–678 (2011).

5. S. Mueller, et al., Individual variability in functional connectivity architecture of the human brain. Neuron 77, 586–595 (2013).

6. C. Gratton, et al., Functional brain networks are dominated by stable group and individual factors, not cognitive or daily variation. Neuron 98, 439-452.e5 (2018).

7. L. Wang, R. E. B. Mruczek, M. J. Arcaro, S. Kastner, Probabilistic maps of visual topography in human cortex. Cereb. Cortex 25, 3911–3931 (2015).

8. M. Carrasco, Visual attention: the past 25 years. Vision Res. 51, 1484–1525 (2011).

9. M. A. Silver, S. Kastner, Topographic maps in human frontal and parietal cortex. Trends Cogn Sci (Regul Ed) 13, 488–495 (2009).

10. K. A. Schneider, M. C. Richter, S. Kastner, Retinotopic organization and functional subdivisions of the human lateral geniculate nucleus: a high-resolution functional magnetic resonance imaging study. J. Neurosci. 24, 8975–8985 (2004).

11. J. Mitchell, C. Bradley, Quality of life in age-related macular degeneration: a review of the literature. Health Qual. Life Outcomes 4, 97 (2006).

12. W. L. Wong, et al., Global prevalence of age-related macular degeneration and disease burden projection for 2020 and 2040: a systematic review and meta-analysis. Lancet Glob. Health 2, e106–16 (2014).

13. S.-H. Cheung, G. E. Legge, Functional and cortical adaptations to central vision loss. Vis. Neurosci. 22, 187–201 (2005).

14. D. C. Fletcher, R. A. Schuchard, Preferred retinal loci relationship to macular scotomas in a low-vision population. Ophthalmology 104, 632–638 (1997).

15. E. S. Finn, et al., Functional connectome fingerprinting: identifying individuals using patterns of brain connectivity. Nat. Neurosci. 18, 1664–1671 (2015).

16. R. M. Braga, R. L. Buckner, Parallel Interdigitated Distributed Networks within the Individual Estimated by Intrinsic Functional Connectivity. Neuron 95, 457-471.e5 (2017).

17. N. Sabbah, et al., Reorganization of early visual cortex functional connectivity following selective peripheral and central visual loss. Sci. Rep. 7, 43223 (2017).

18. L. L. Fleming, et al., Impact of Deprivation and Preferential Usage on Functional Connectivity Between Early Visual Cortex and Category-Selective Visual Regions. Hum. Brain Mapp. 45, e70064 (2024).

19. A. M. Larson, L. C. Loschky, The contributions of central versus peripheral vision to scene gist recognition. J. Vis. 9, 6. 1–16 (2009).

20. D. G. Pelli, et al., Crowding and eccentricity determine reading rate. J. Vis. 7, 20. 1–36 (2007).

21. S.-A. Yoo, S. C. Chong, Eccentricity biases of object categories are evident in visual working memory. Vis. cogn. 20, 233–243 (2012).

22. A. Trouilloud, et al., Rapid scene categorization: From coarse peripheral vision to fine central vision. Vision Res. 170, 60–72 (2020).

23. L. Zhaoping, Feedback from higher to lower visual areas for visual recognition may be weaker in the periphery: Glimpses from the perception of brief dichoptic stimuli. Vision Res. 136, 32–49 (2017).

24. J. C. Griffis, A. S. Elkhetali, W. K. Burge, R. H. Chen, K. M. Visscher, Retinotopic patterns of background connectivity between V1 and fronto-parietal cortex are modulated by task demands. Front. Hum. Neurosci. 9, 338 (2015).

25. S. A. Sims, P. Demirayak, S. Cedotal, K. M. Visscher, Frontal cortical regions associated with attention connect more strongly to central than peripheral V1. Neuroimage 238, 118246 (2021).

26. L. Zhaoping, J. Ackermann, Reversed Depth in Anticorrelated Random-Dot Stereograms and the Central-Peripheral Difference in Visual Inference. Perception, 301006618758571 (2018).

27. L. Zhaoping, The Flip Tilt Illusion: Visible in Peripheral Vision as Predicted by the Central-Peripheral Dichotomy. Iperception 11, 2041669520938408 (2020).

28. J. M. Tangen, S. C. Murphy, M. B. Thompson, Flashed face distortion effect: grotesque faces from relative spaces. Perception 40, 628–630 (2011).

29. D. M. Little, K. R. Thulborn, J. P. Szlyk, An FMRI study of saccadic and smooth-pursuit eye movement control in patients with age-related macular degeneration. Invest. Ophthalmol. Vis. Sci. 49, 1728–1735 (2008).

30. C. Baldassano, D. M. Beck, L. Fei-Fei, Human-Object Interactions Are More than the Sum of Their Parts. Cereb. Cortex 27, 2276–2288 (2017).

31. K. V. Haak, A. B. Morland, S. A. Engel, Plasticity, and its limits, in adult human primary visual cortex. Multisens. Res. 28, 297–307 (2015).

32. B. A. Wandell, S. M. Smirnakis, Plasticity and stability of visual field maps in adult primary visual cortex. Nat. Rev. Neurosci. 10, 873–884 (2009).

33. C. D. Gilbert, W. Li, Adult visual cortical plasticity. Neuron 75, 250–264 (2012).

34. O. H. Butt, N. C. Benson, R. Datta, G. K. Aguirre, The fine-scale functional correlation of striate cortex in sighted and blind people. J. Neurosci. 33, 16209–16219 (2013).

35. H. A. Baseler, et al., Large-scale remapping of visual cortex is absent in adult humans with macular degeneration. Nat. Neurosci. 14, 649–655 (2011).

36. S. M. Smirnakis, et al., Lack of long-term cortical reorganization after macaque retinal lesions. Nature 435, 300–307 (2005).

37. Y. Masuda, S. O. Dumoulin, S. Nakadomari, B. A. Wandell, V1 projection zone signals in human macular degeneration depend on task, not stimulus. Cereb. Cortex 18, 2483–2493 (2008).

38. H. Abe, et al., Adult cortical plasticity studied with chronically implanted electrode arrays. J. Neurosci. 35, 2778–2790 (2015).

39. K. V. Haak, A. B. Morland, G. S. Rubin, F. W. Cornelissen, Preserved retinotopic brain connectivity in macular degeneration. Ophthalmic Physiol. Opt. 36, 335–343 (2016).

40. C. I. Baker, E. Peli, N. Knouf, N. G. Kanwisher, Reorganization of visual processing in macular degeneration. J. Neurosci. 25, 614–618 (2005).

41. D. D. Dilks, C. I. Baker, E. Peli, N. Kanwisher, Reorganization of visual processing in macular degeneration is not specific to the “preferred retinal locus”. J. Neurosci. 29, 2768–2773 (2009).

42. Y. Masuda, et al., V1 projection zone signals in human macular degeneration depend on task despite absence of visual stimulus. Curr. Biol. 31, 406-412.e3 (2021).

43. A. S. Bock, et al., Resting-State Retinotopic Organization in the Absence of Retinal Input and Visual Experience. J. Neurosci. 35, 12366–12382 (2015).

44. Fine, J.-M. Park, Blindness and human brain plasticity. Annu. Rev. Vis. Sci. 4, 337–356 (2018).

45. S. Sen, et al., The role of visual experience in individual differences of brain connectivity. J. Neurosci. 42, 5070–5084 (2022).

46. E. Striem-Amit, et al., Functional connectivity of visual cortex in the blind follows retinotopic organization principles. Brain 138, 1679–1695 (2015).

47. C. D. Gilbert, W. Li, Top-down influences on visual processing. Nat. Rev. Neurosci. 14, 350–363 (2013).

48. T. Plank, et al., Cortical thickness related to compensatory viewing strategies in patients with macular degeneration. Front. Neurosci. 15, 718737 (2021).

49. W. K. Burge, et al., Cortical thickness in human V1 associated with central vision loss. Sci. Rep. 6, 23268 (2016).

50. M. K. Defenderfer, et al., Cortical plasticity in central vision loss: Cortical thickness and neurite structure. Hum. Brain Mapp. 44, 4120–4135 (2023).

51. M. Welvaert, Y. Rosseel, On the definition of signal-to-noise ratio and contrast-to-noise ratio for FMRI data. PLoS ONE 8, e77089 (2013).

52. S. A. M. Stevelink, E. M. Malcolm, N. T. Fear, Visual impairment, coping strategies and impact on daily life: a qualitative study among working-age UK ex-service personnel. BMC Public Health 15, 1118 (2015).

53. R. L. Buckner, F. M. Krienen, B. T. T. Yeo, Opportunities and limitations of intrinsic functional connectivity MRI. Nat. Neurosci. 16, 832–837 (2013).

54. R. Patriat, et al., The effect of resting condition on resting-state fMRI reliability and consistency: a comparison between resting with eyes open, closed, and fixated. Neuroimage 78, 463–473 (2013).

55. A. B. Waites, A. Stanislavsky, D. F. Abbott, G. D. Jackson, Effect of prior cognitive state on resting state networks measured with functional connectivity. Hum. Brain Mapp. 24, 59–68 (2005).

56. R. J. Barry, et al., Caffeine effects on resting-state arousal. Clin. Neurophysiol. 116, 2693–2700 (2005).

57. C. W. Wong, V. Olafsson, O. Tal, T. T. Liu, Anti-correlated networks, global signal regression, and the effects of caffeine in resting-state functional MRI. Neuroimage 63, 356–364 (2012).

58. G. Deco, P. Hagmann, A. G. Hudetz, G. Tononi, Modeling resting-state functional networks when the cortex falls asleep: local and global changes. Cereb. Cortex 24, 3180–3194 (2014).

59. L. Tarita-Nistor, E. G. González, S. N. Markowitz, M. J. Steinbach, Plasticity of fixation in patients with central vision loss. Vis. Neurosci. 26, 487–494 (2009).

60. S. Sunness, C. A. Applegate, D. Haselwood, G. S. Rubin, Fixation patterns and reading rates in eyes with central scotomas from advanced atrophic age-related macular degeneration and Stargardt disease. Ophthalmology 103, 1458–1466 (1996).

61. E. Guez, J. F. Le Gargasson, F. Rigaudiere, J. K. O’Regan, Is there a systematic location for the pseudo-fovea in patients with central scotoma? Vision Res. 33, 1271–1279 (1993).

62. U. L. Nilsson, C. Frennesson, S. E. Nilsson, Patients with AMD and a large absolute central scotoma can be trained successfully to use eccentric viewing, as demonstrated in a scanning laser ophthalmoscope. Vision Res. 43, 1777–1787 (2003).

63. U. L. Nilsson, Visual rehabilitation with and without educational training in the use of optical aids and residual vision. A prospective study of patients with advanced age-related macular degeneration. Clinical Vision Sciences 6, 3–10 (1990).

64. Carrasco, A. M. Giordano, B. McElree, Temporal performance fields: visual and attentional factors. Vision Res. 44, 1351–1365 (2004).

65. Carrasco, C. P. Talgar, E. L. Cameron, Characterizing visual performance fields: effects of transient covert attention, spatial frequency, eccentricity, task and set size. Spat. Vis. 15, 61–75 (2001).

66. E. A. Piazza, M. A. Silver, Persistent hemispheric differences in the perceptual selection of spatial frequencies. J. Cogn. Neurosci. 26, 2021–2027 (2014).

67. A. K. M. R. Karim, H. Kojima, The what and why of perceptual asymmetries in the visual domain. Adv. Cogn. Psychol. 6, 103–115 (2010).

68. J. Intriligator, P. Cavanagh, The spatial resolution of visual attention. Cogn. Psychol. 43, 171–216 (2001).

69. J. Abrams, A. Nizam, M. Carrasco, Isoeccentric locations are not equivalent: the extent of the vertical meridian asymmetry. Vision Res. 52, 70–78 (2012).

70. A. Barbot, S. Xue, M. Carrasco, Asymmetries in visual acuity around the visual field. J. Vis. 21, 2 (2021).

71. M. Himmelberg, J. Winawer, M. Carrasco, Stimulus-dependent contrast sensitivity asymmetries around the visual field. J. Vis. 20, 18 (2020).

72. S. Fuller, R. Z. Rodriguez, M. Carrasco, Apparent contrast differs across the vertical meridian: visual and attentional factors. J. Vis. 8, 16. 1–16 (2008).

73. Rubin, K. Nakayama, R. Shapley, Enhanced perception of illusory contours in the lower versus upper visual hemifields. Science 271, 651–653 (1996).

74. K. Visscher, P. Stewart, Macular Degeneration and Plasticity (MDP): Connectomes in Human Diseases (2024) (January 3, 2025).

75. F. L. Ferris, A. Kassoff, G. H. Bresnick, I. Bailey, New visual acuity charts for clinical research. Am. J. Ophthalmol. 94, 91–96 (1982).

76. M. F. Glasser, et al., The Human Connectome Project’s neuroimaging approach. Nat. Neurosci. 19, 1175–1187 (2016).

77. K. J. Gorgolewski, et al., The brain imaging data structure, a format for organizing and describing outputs of neuroimaging experiments. Sci. Data 3, 160044 (2016).

78. O. Esteban, et al., MRIQC: Advancing the automatic prediction of image quality in MRI from unseen sites. PLoS ONE 12, e0184661 (2017).

79. O. Esteban, et al., fMRIPrep: a robust preprocessing pipeline for functional MRI. Nat. Methods 16, 111–116 (2019).

80. K. J. Gorgolewski, et al., Nipype: a flexible, lightweight and extensible neuroimaging data processing framework in python. Front. Neuroinformatics 5, 13 (2011).

81. A. M. Dale, B. Fischl, M. I. Sereno, Cortical surface-based analysis. I. Segmentation and surface reconstruction. Neuroimage 9, 179–194 (1999).

82. B. B. Avants, C. L. Epstein, M. Grossman, J. C. Gee, Symmetric diffeomorphic image registration with cross-correlation: evaluating automated labeling of elderly and neurodegenerative brain. Med. Image Anal. 12, 26–41 (2008).

83. Y. Zhang, M. Brady, S. Smith, Segmentation of brain MR images through a hidden Markov random field model and the expectation-maximization algorithm. IEEE Trans. Med. Imaging 20, 45–57 (2001).

84. D. N. Greve, B. Fischl, Accurate and robust brain image alignment using boundary-based registration. Neuroimage 48, 63–72 (2009).

85. M. Jenkinson, P. Bannister, M. Brady, S. Smith, Improved optimization for the robust and accurate linear registration and motion correction of brain images. Neuroimage 17, 825–841 (2002).

86. R. W. Cox, J. S. Hyde, Software tools for analysis and visualization of fMRI data. NMR in Biomedicine (1997).

87. J. D. Power, et al., Methods to detect, characterize, and remove motion artifact in resting state fMRI. Neuroimage 84, 320–341 (2014).

88. Y. Behzadi, K. Restom, J. Liau, T. T. Liu, A component based noise correction method (CompCor) for BOLD and perfusion based fMRI. Neuroimage 37, 90–101 (2007).

89. C. Lanczos, Evaluation of noisy data. Journal of the Society for Industrial and Applied Mathematics Series B Numerical Analysis 1, 76–85 (1964).

90. J. D. Power, K. A. Barnes, A. Z. Snyder, B. L. Schlaggar, S. E. Petersen, Spurious but systematic correlations in functional connectivity MRI networks arise from subject motion. Neuroimage 59, 2142–2154 (2012).

91. K. R. A. Van Dijk, M. R. Sabuncu, R. L. Buckner, The influence of head motion on intrinsic functional connectivity MRI. Neuroimage 59, 431–438 (2012).

92. R. Ciric, et al., Mitigating head motion artifact in functional connectivity MRI. Nat. Protoc. 13, 2801–2826 (2018).

93. E. W. Dickie, et al., Ciftify: A framework for surface-based analysis of legacy MR acquisitions. Neuroimage 197, 818–826 (2019).

94. D. S. Marcus, et al., Informatics and data mining tools and strategies for the human connectome project. Front. Neuroinformatics 5, 4 (2011).

95. N. C. Benson, J. Winawer, Bayesian analysis of retinotopic maps. eLife 7 (2018).

96. M. Defenderfer, P. Demirayak, K. M. Visscher, A method for mapping retinal images in early visual cortical areas. Neuroimage 245, 118737 (2021).

97. R. M. Steinman, Effect of target size, luminance, and color on monocular fixation*. J. Opt. Soc. Am. 55, 1158 (1965).

98. A. M. Winkler, G. R. Ridgway, M. A. Webster, S. M. Smith, T. E. Nichols, Permutation inference for the general linear model. Neuroimage 92, 381–397 (2014).

99. M. J. Anderson, J. Robinson, Permutation Tests for Linear Models. Aust NZ J Stat 43, 75–88 (2001).

